# Unsuspected transcriptional regulations during rice defense response revealed by a toolbox of marker genes for rapid and extensive analysis of expression changes upon various environments

**DOI:** 10.1101/2022.12.14.520374

**Authors:** Pélissier Rémi, Brousse Andy, Ramamonjisoa Anjara, Ducasse Aurélie, Ballini Elsa, Jean-Benoit Morel

## Abstract

Since rice (*Oryza sativa*) is an important crop and the most advanced model for monocotyledonous species, acceding to its physiological status is important for many fundamental and applied purposes. Although this physiological status can be obtained by measuring the transcriptional regulation of marker genes, the tools to perform such analysis are often too expensive, non flexible or time consuming. Here we manually selected 96 genes considered as biomarkers of important processes taking place in rice leaves based on literature analysis. We monitored their transcriptional regulation under several treatments (disease, phytohormone inoculation, abiotic stress…) using Fluidigm method that allows to perform ~10 000 RT-QPCR reactions in one single run. This technique allowed us to verify a large part of known regulations but also to identify new, unsuspected regulations. Together, our set of genes, coupled to our data analysis protocol with Fluidigm brings a new opportunity to have a fast and reasonably cheap access to the physiological status of rice leaves in a high number of samples.

## Introduction

Monitoring the physiological status of plants is important for several purposes, for instance to understand how plants react to stimuli when studying stress biology. It also becomes more and more necessary to monitor plant molecular status in the context of agro-ecological practices such as growth stimulation or biocontrol product development for which a mechanism of action is often required. In both cases, biologists are facing a challenge for the study of response to multiple combined factors since it generates a large set of samples to be analyzed, with cost limitations. For instance, the analysis of the combination of 2 treatments (and their corresponding controls) in 4 time points across 6 genotypes in 4 replicates requires the analysis of 384 samples.

The need for a rapid and efficient assessment of plant status, along with the potential to simultaneously quantify multiple traits on a large number of individuals plant over multiple stress and time periods has led to the development of new technologies for plant science (Galieni et al., 2021). For instance, in the last decade, a large variety of imaging techniques (visible, spectroscopy, fluorescence and infrared imaging) has been a powerful tool to collect data of complex traits related to the growth, yield and plant adaptation to different treatments or stress (Li et al., 2014).

At the cellular level, the development of new high throughput methods for the analysis of biological molecules (e.g “Omic”, Hennig, 2007) has enabled screening for molecular biomarkers (measurable biological molecules that are characteristic for a specific physiological status). Molecular biomarkers can be identified at different scales in the cell by analyzing the proteome, metabolome or the analysis of lipids for instance (Riedmaier and Pfaffl, 2013). Since protein/enzyme production initially relies on gene transcription, transcription of marker genes can be used to monitor of the physiological status of a tissue (Biswas et al., 2017). While biomarkers have long been identified under specific conditions without testing other conditions, nowadays marker genes can be identified *in silico* from the analysis of a large number of conditions (Chen et al., 2003, 2002; Cohen and Leach, 2019; Miao et al., 2017). However, in most studies the *in vivo* expression of candidate marker genes has been done on a limited set conditions (Riedmaier and Pfaffl, 2013). For instance, we previously published a set of 24 genes useful to characterize defense response in rice (Delteil et al., 2012).

Although transcription and metabolic pathways are not fully correlated, RNA quantity of markers genes can be used as a proxy (Schwahn and Nikoloski, 2018). In that sense, quantitative RT-PCR has clearly opened new perspectives in plant biology. Recent advances in high-throughput technologies to survey RNA, especially microarray profiling and RNA sequencing (RNA-seq), have revolutionized the discipline and enabled the study of gene expression at the level of whole transcriptome rather than individual transcripts, typically targeted by Northern blot or Real Time quantitative PCR (Martin et al., 2016). Traditional techniques such RT-QPCR are precise and low cost, but they only focus on expression of one gene and remain time consuming to access to multiple gene expression (Caldana et al., 2007). On the contrary, highly parallel profiling technologies (Omics) such as RNAseq allow research to comprehensively analyze cellular status (Mantione et al., 2014) but they remain costly and demanding for bioinformatics skills and computational effort (Goralski et al., 2016). Accessing rapidly and cost-effectively to medium-scale expression of genes representative of different pathways for research purposes remains a challenge.

The Fluidigm technology (www.fluidigm.com), a nanofluidic automated real-time PCR system that relies on microfluidic technology with dynamic arrays of integrated fluidic circuits, fills the gap between RNAseq and traditional RT-QPCR analysis. In this method, typical chip format allows for 9 216 RT-QPCR simultaneous reactions (96.96 chip format; 96 samples × 96 assays) in a single qPCR run. This makes Fluidigm more cost-effective compared to other Omics technologies, more flexible than microarray and less time-consuming than RT-QPCR.

In this study, based on literature analysis, we selected a set of 96 genes of rice whose transcription is regulated in leaves by biotic, abiotic stresses or that are known markers of major hormonal, primary or secondary metabolisms. The regulation of the expression of these genes was assessed under different biotic, abiotic stresses and phytohormone treatment using Fluidigm technology. Together Fluidigm and our set of genes brings a new method that is cheap, rapid and reliable to access to the global transcriptomic status of rice leaves, in particular under biotic stresses.

## Material and methods

### Experimental setup, plant material and growth conditions

To validate *in vivo* our set of marker genes identified from bibliography analysis, we ran one validation experiment. We treated rice with different phytohormones or abiotic/biotic stress separately with their corresponding controls treatment. We used the same rice genotype Nipponbare of *Oryza sativa* sp temperate japonica and the same number of replications for all experiments. All experiments were conducted in greenhouse with the same growing condition. Five plants of Nipponbare were grown in plastic pots (9×9×9,5 cm) filled with substrate (58% blond peat, 18% coconut powder, 10% perlite, 9% volcanic sand, 5% clay) supplemented with 3,5g/L of fertilization (Basacote Native 6M, NPK 14-3-19) (except for nitrogen stress). Pots were normally watered each day (except for Drought treatment) and grown under 16h artificial light (55000 lumen) at 27°-23°C.

### Phytohormone treatments

After 3 weeks of growth, rice leaves were sprayed with individual phytohormones: Abscisic acid (ABA), Methyl jasmonate as jasmonic acid substitute (JA), Ethephon as Ethylene substitute (ET), auxin (IAA), Kinetin as cytokinin substitute (CK) and salicylic acid (SA). For each phytohormone, three solutions were prepared and sprayed on rice leaves: two at different concentrations based on literature and one with the solvent solution without phytohormone used as mock treatment (**Supplementary Table.S1)**. For each treatment, 30mL of solution was sprayed on 6 pots of five plants each (1mL/plant). Leaf tissues for RNA extraction were collected 6 hours after treatment. Plants treated with volatile phytohormones (abscisic acid, ethephon or methyl jasmonate) as well as the corresponding controls were placed under airtight plastic cages during incubation to prevent evaporation and diffusion of the corresponding chemicals.

### Stress treatments

#### Drought stress

For drought stress, we used a protocol adapted from (Bidzinski et al., 2016). Briefly, plants were cultivated as described above with the same amount of fertilization and watered each day. At day 19, trays of both mock treatment and water stress were flooded with 2 to 3 cm of water. At day 23, drought stress was imposed by removing water to 6 pots and stopping watering for 5 days before tissue harvest. The 6 mock pots remained inundated by water during these 5 days (**Supplementary Figure S1 a**).

#### Nitrogen stress (depletion)

For nitrogen stress, rice plants were sown in pots and cultivated as described above but without any fertilization added previously in the soil. At days 7 and 14, all pots of mock and nitrogen stressed treatment were fertilized with a N1 liquid solution containing 50%NH4+/50%NO3-(40 mg/L) of nitrogen and all other nutrients needed for rice growth as described in (Ballini et al., 2013). At day 21, mock treated plants were fertilized with the N1 solution and the nitrogen stress plants were fertilized with the N0 solution containing all nutrients except nitrogen. Six pots were used for each mock and nitrogen stress treatment and tissue was performed at day 28 (**Supplementary Figure S1 b**).

#### Magnaporthe oryzae inoculation

For pathogen inoculation, we used the multivirulent *Guy11* strain (Gallet et al., 2015) of hemibiotrophic fungal pathogen that was grown for 10 days on rice flour agar medium (20 g of rice flour, 15 g of agar, 2.5 g of yeast extract and 1 L of distilled water) under fluorescent light (12 h/day) at 26°C. We harvested conidia by flooding the plate with 5 ml of sterile distilled water. Suspensions (with 0.1% gelatin) of 30 mL of a 50000 (concentration 1) or 100000 (concentration 2) conidia per ml were sprayed on each tray containing 6 pots on three-week old plants as described in Berruyer *et al*., 2003. Besides inoculation with spores, a spray with the inoculation solution (0.1% gelatin) without spore was also used on 6 other pots as a mock treatment. After inoculation, rice plants were incubated for 16 h in a controlled climatic chamber at 25°C with 95% relative humidity and later returned to normal growth conditions. Tissue harvest was performed 24h after inoculation when defenses are usually first expressed (Delteil et al., 2012).

### Gene selection and primer design

A systematic literature review was conducted across three databases PubMed, Google scholar, and Science direct to identify gene biomarkers of selected processes. If available, we preferably selected studies showing gene transcriptomic regulation (by RT-QPCR, Microarray or RNA-seq) in rice young seedling leaf related to a targeted process (**Supplementary Table.S2**). When we could not find studies reporting transcriptomic regulation of gene in one targeted process, we selected studies showing the implication of genes using mutant approaches.

If available, we selected already designed PCR primers based on our bibliography analysis. However if no primers were already published for the gene targeted, we designed our own primers. The design of primers followed a set of stringent criteria, as generally suggested in qRT-PCR protocols (e.g. PrimerExpress Software v2.0 Application Manual, Applied Biosystems). To minimize the risk of amplifying contaminating genomic DNA, primers spanning at least one exon-exon junction, or annealing to different exons, were designed when possible. The specificity of each primer was confirmed by comparing its sequence with all predicted rice coding sequences (CDS) using the Primer3Plus software to ensure that at least one primer of each pair targets a unique site within the set of predicted rice CDS. Primers specificity and amplicon length were checked for each pair of primers using MFEprimer3.1 and verification of Hairpin and homo/heterodimerization was checked with idtdna OligoAnalyzer.

### RNA extraction and transcriptomic assay

For each treatment at each concentration (Mock, concentration 1, concentration 2) we used at least 6 pots 5 plants each from which me built 6 RNA as follows: each RNA represented a mix of the middle part of 3 last developed leaves randomly selected from 30 plants to limit position effect on gene expression. This represented 144 RNA samples extracted and analyzed in this study.

For RNA extraction, we used protocols described in Delteil *et al*. (2012). Briefly, frozen leaf tissues were ground in liquid nitrogen. Approximately 500 mg of powder was treated with 1 mL of TRIZOL (Invitrogen, Carlsbad, CA, USA). RNA samples (5μg) were denatured for 5 min at 65°C with oligo(dT) 18 (3.5 mM) and deoxynucleoside triphosphate (dNTP) (1.5 mM). They were later subjected to reverse transcription for 45 min at 37°C with 200 U of reverse transcriptase M-MLV (Promega, Madison, WI, USA) in the appropriate buffer.

For classical RT-QPCR analysis (comparaison with fluidigm data and primer optimization), two microliters of cDNA (dilution 1:10) were used for amplification. RT-QPCR mixtures contained PCR buffer, dNTP (0.25 mM), MgCl2 (2.5 mM), forward and reverse primers (final concentration of 150, 300 or 600 nM), 1 U of HotGoldStar polymerase and SYBR Green PCR mix as per the manufacturer’s recommendations (Eurogentec, Seraing, Belgium). Amplification was performed as follows: 95°C for 10 min; 40 cycles of 95°C for 15 s, 62°C for 1 min and 72°C for 30 s; finally, 95°C for 1 min and 55 °C for 30 s. The RT-QPCRs were performed using a LightCycler480 machine (Roche; Mannheim, Germany) and data were extracted using LightCycler480 software. The amount of plant RNA in each sample was normalized using actin (Os03g50890) for rice as internal control. The calculation of gene expression was performed using the measured efficiency for each primer pair as described in Vergne *et al*., 2007.

For Fluidigm gene expression analysis, two μL of cDNA samples were pre-amplified following the “Pre-amplification of cDNA for Gene Expression with delta Gene Assays” protocol provided by the manufacturer (BIOMARK HD, Fluidigm). The reactions were cleaned up using Exonuclease I (Exo I, 4U/uL). Gene expression analysis was performed with the 96.96 IFC Machine (San Francisco, CA) using the Delta Gene Assays and he protocol provided by Fluidigm on 1/10 dilluted pre-amplified samples (5 μL gene assay mix and sample assays used for running the plate). Amplification was performed as follows: 95°C for 1 min; 30 cycles of 96°C for 5 s and 60°C for 1 min; finally, 95°C for 1 min. Gene expression was measured for 94 genes identified in the literature. Out of the 92 genes initially selected that were known to be transcriptionally regulated in the different targeted processes (**Table S2**), only 88 were analyzed (**Table S3**). Baseline correction (using linear derivative) and assessment of cycle threshold (Ct) values were performed by the BIOMARK HD software (Fluidigm). Fluidigm gene expression was calculated by the 2 ΔΔCt method (Livak and Schmittgen, 2001) using the 3 most stable references genes EF1a, EF4a and UBQ5 with the R package *“fluidigr”*. Genes for which expression was not detected in the entire data sample have been deleted from the analysis.

### Data analysis and statistics

All analyses were made using R (R Core Team, 2019). For clustering analysis, a Principal Component Analysis (PCA) was performed using the R package *“FactomineR”*. Clustering of gene expression and heatmap visualization were done with the R package *“gplots.heatmap”* and *“ggplot2”*.

For statistical analysis, we first identified significant differences between mock and treated plants in the gene expression analysis. Analysis of distribution of expression value (2–ΔΔCt) revealed that at least one of the distribution was not normal, thus we evaluated differences between conditions using non-parametric Wilcoxon’s tests (with a Bonferroni correction). Second, in order to identify marker genes, for each gene we compared fold change values between treatments. Fold change were calculated by dividing values of the expression of each gene for each treatment by the mean of the gene in the corresponding mock condition. After that step, a linear model was applied for each gene where the log transformed fold change was a function of the condition (treatment x concentration). Log transformation was used to correct for normality and homocedasticity. The impact of each condition on gene fold change was tested by ANOVA followed by a Tukey HSD test (Function *glht* from the package *multcomp*).

## Results

### Building of an expert rice gene platform for quantitative PCR

We conducted a large bibliography analysis and identified genes known to be transcriptionally regulated in the targeted process (see Methods and Table. S1). Genes were individually selected carefully based on the analysis of literature following the following guidelines: i) shown evidence of transcriptomic regulation in leaf and ii) demonstrated transcriptomic regulation by external stimuli (environmental stress / treatment …) or gene implicated in the response, homeostasis or signaling of the targeted process **(Figure 1).**

**Figure.1.**
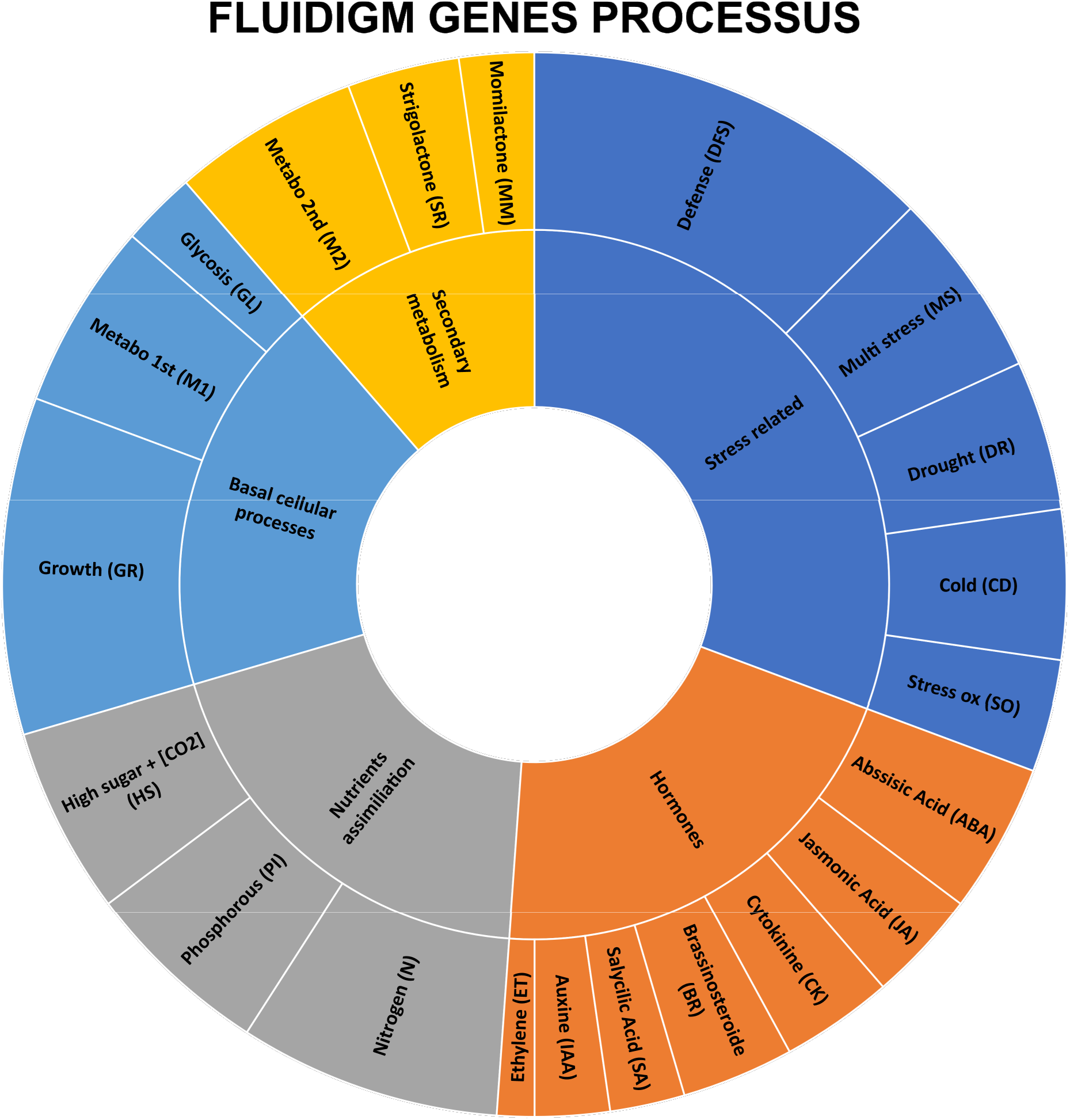
Distribution of physiological processes represented in the gene expression platform. Genes were selected based on bibliography analysis of genes transcriptionally regulated in rice leaf seedlings. Complete list of genes, regulation and reference is available in **Table.S1.** Gene were selected to cover major physiological processes such as:

- **Homonal**: Abscissic Acid (**ABA**), Salycilic acid (**SA**), Jasmonic acid (**JA**), Auxin (**IAA**), Cytokinine (**CK**), Ethylene (**ET**);
- **Stress related**: Multi stress (**MS**), response to abiotic stress (Cold (**CD**), drought (**DR**), oxidative stress (**SO**)) or biotic stress: defense pathways (**DFS**).
- **Secondary metabolism**: Secondary metabolism synthesis (**M2**) (flavonoid, phyotalexine), Momilactone synthesis (**MM**), strigolactone (**SR**).
- **Basal cellular processes** sucha as primary metabolism:Krebs cycle (**M1**), glycolysis (**GL**) and growth (**GR**).
- **Nutrient assimilation**: nitrogen (**N**), phosphorous (**PI**)) Sugar and CO_2_ concentration (**HS**)).

In total, out of these 150 genes initially identified (data not shown), we selected 92 genes known to be expressed and transcriptionally regulated in rice leaf and 4 reference genes commonly used in rice transcriptome analysis were added **(Supplementary Table.S2).** Biological processes were selected to target major rice hormonal pathways (Abscissic Acid, Salycilic acid, Jasmonic acid, Auxin, Cytokinine, Ethylene, Gibberelin, Brassinoteroide and Strigolactone). Genes were also selected to represent basal cellular processes (Krebs cycle / primary metabolism, glycolysis, growth), secondary metabolism (flavonoid, phyotalexine), momilactone production, response to abiotic / biotic stress (Cold, drought, oxidative stress, defense) or nutrient assimilation (nitrogen, phosphorous, Sugar/CO_2_ concentration). There were on average 2-4 marker genes per process, thus maximizing the chances to detect transcriptional changes for each process.

### Analysis of gene regulation under treatments

Among the 96 genes selected and optimized for classical RT-QPCR, four (OsAOS2, OsKS4, IPA1 and OsLipoxy) were not detected or very weakly expressed with the Fluidigm technology and were removed from following analysis. Therefore we could analyze the regulation of a set of 88 marker genes and 4 constitutive control genes. The transcriptional regulation of these 88 marker genes was analyzed under several individual treatments (phytohormone exogenous application, *M. oryzae* inoculation, drought stress and nitrogen stress). For most treatments, two different concentrations (except drought and nitrogen stress) were tested in addition to the mock treatment representing the control condition.

To test if quantification of mRNA was correlated between Fluidigm technology and classical QRT-PCR, we randomly picked 5 genes in 4 random treatments and checked if their expression correlated with the one obtained from Fluidigm analysis (**Figure S2**). A significative Pearson correlation (r^2^ = 0.45, p < 0.001) was detected between QRT-PCR and Fluidigm analysis.

Given our RNA set and the available treatments tested, published transcriptional regulation could be expected for ~50% of the selected genes (47 genes transcriptionally regulated by hormone treatments, drought, nitrogen and pathogen stresses). In contrast, as we did not have the capacity to produce differential samples for some pathways (e.g. primary metabolism), there were 41 genes for which we had no particular expectation in terms of transcriptional regulation in our RNA set.

To evaluate the regulation of each gene by the different treatments, we performed a statistical comparison of the expression of each gene between the mock and treated condition. We found 161 significative differences (Wilcoxon test with a Bonferoni correction; reported on the **Figure 2** by a star symbols) confirming 30/47 (~ 64%) of the expected regulations (**Figure 3**).

**Figure 2.**
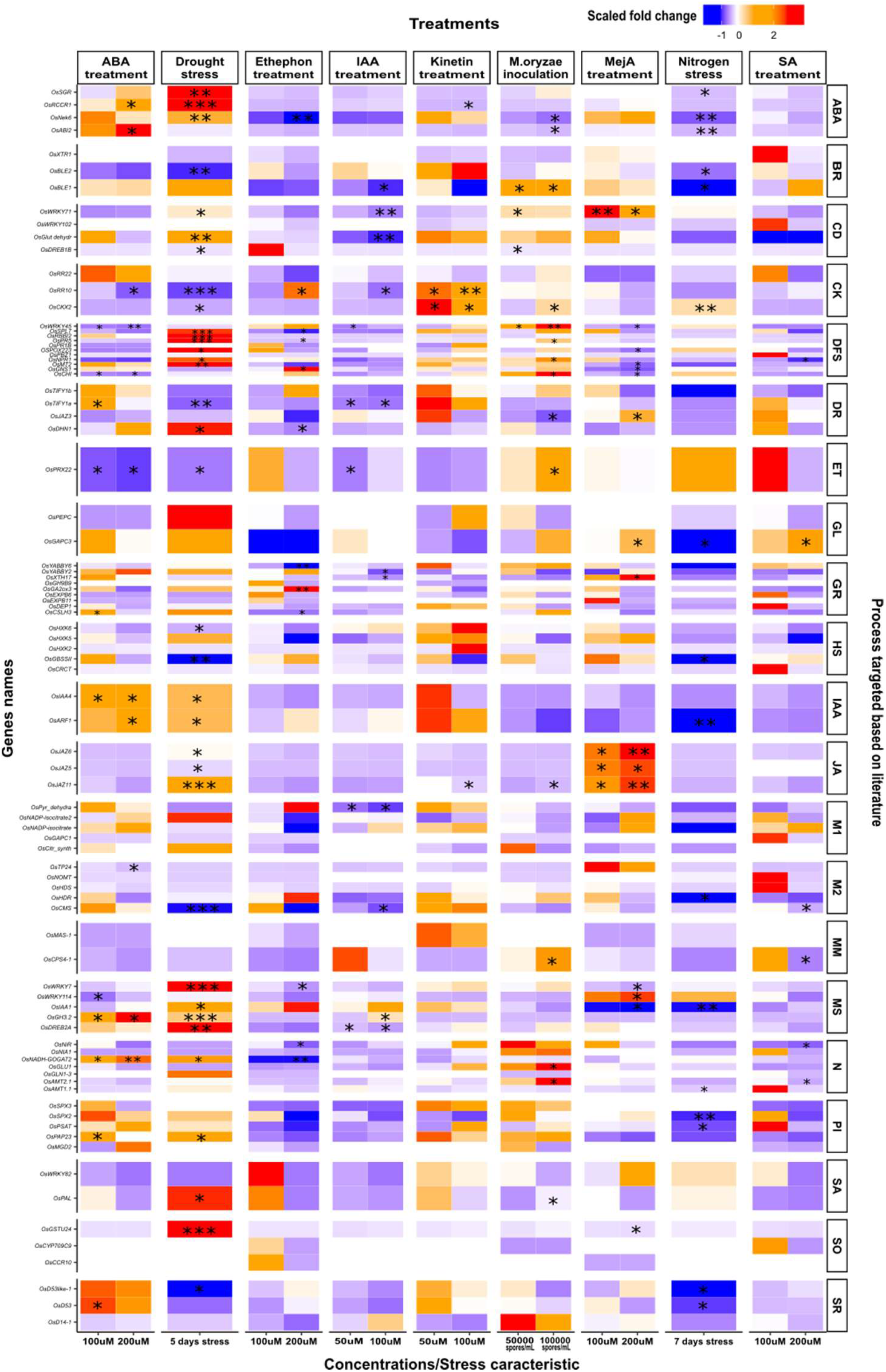
Heatmap of gene expression in response to nine treatments. Gene expression was measured in leaves of rice plants grown and treated as described in the Methods section. The expression of each gene measured by Fluidigm was normalized using 3 reference genes (UBQ5, EF1a, EF4a, 2–ΔΔCt method). Fold change were calculated by dividing the mean of gene expression in treated condition by the mean of the gene in mock condition for each treatment / gene (see Methods). Colors represent the scaled fold change values compared to mock (Blue = downregulated, white = no change, red = upregulated). The statistical differences, as estimated by Wilcoxon tests corrected by a Bonferroni correction, between treated and mock conditions are shown (*: p < 0,05 ; **: p < 0,01, ***: p < 0,001).

**Figure 3.**
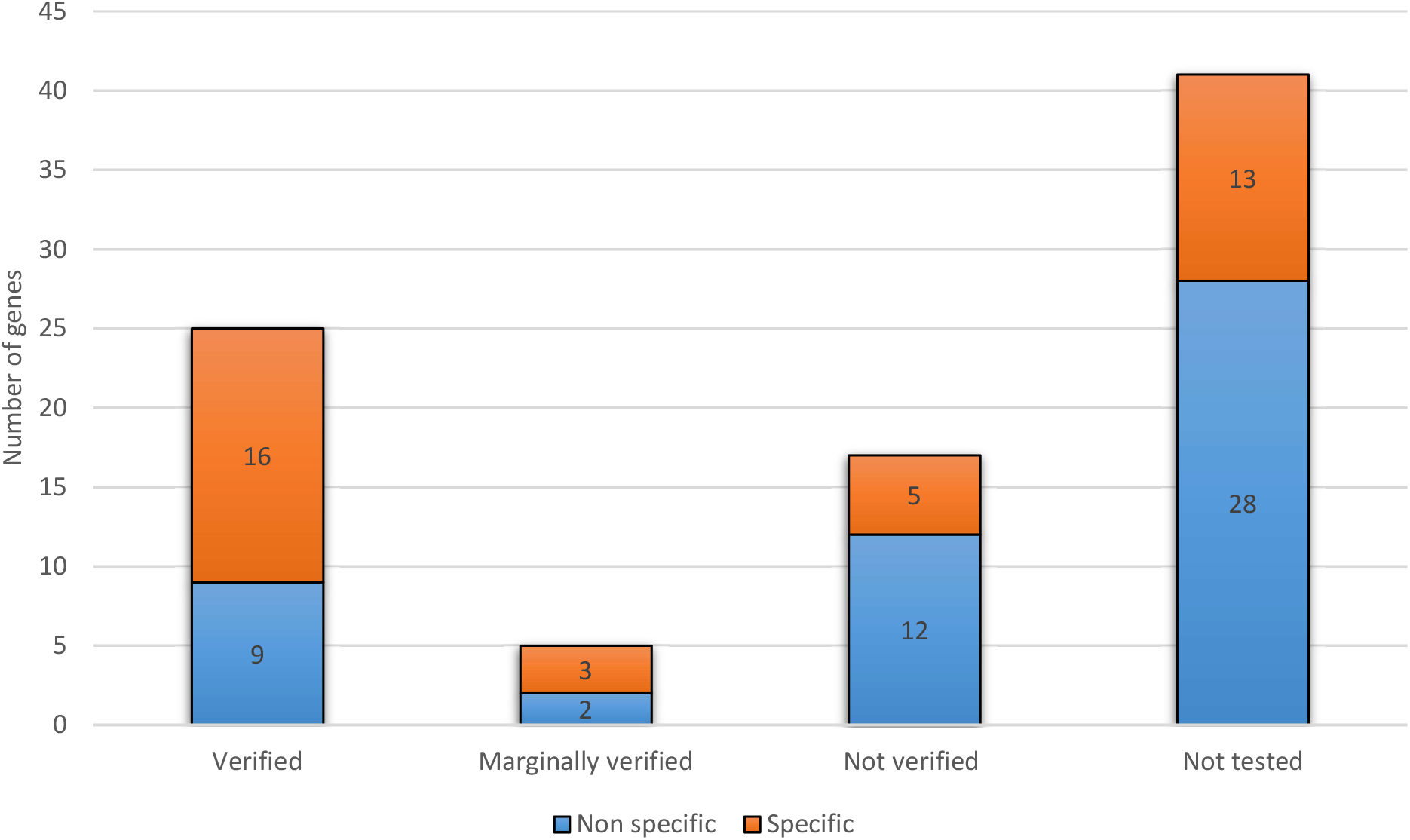
Gene regulation analysis results compared to literature information. Analysis of the gene regulation under treatment or stress in our study compared to the original bibliography data used for selecting the genes. The number of genes presenting the expected regulation were noted as “verified” and the gene presenting a tendency towards the expected regulation were noted as “marginally verified”. Gene regulations that were not similar to the literature information were noted as “Not verified”. Gene whose regulation could not be verified by our treatment (e.g cold, momilactone or phosphorous content responsive gene) were noted as “Not tested”. Gene that were mostly regulated by one treatment or two treatment were noted as specific (orange bars) while the others (induced by multiple treatment the same ways or not modified by any treatment for instance) were considered as not specific (blue bars).

Next, to identify marker genes specifically responding to only one or two treatments, we compared fold changes obtained from each gene in each treatment using a Tukey HSD (see Methods). When considering specificity of expression in this manner in our dataset, we found 37 genes specifically responding to one or two treatments out of the nine treatments covered by our RNA set. Beyond the genes considered as markers of one of our 9 treatment, we detected at least one marker gene for each treatment (OsABI2 for ABA; OsCKX2 for CK; OsCHI for defense, DREB2A/WRKY7 for Drought; OsJAZ5/OsJAZ11 for JA and OsPBZ1 for SA treatment). However, we did not identify specific marker genes for Auxin and Ethylene treatment or Nitrogen stress (**Figure 3**; **Supplementary Table.S3**).

## Discussion

Despite the increasing availability of Omics tools and methods, especially for transcriptomic analysis (e.g RNAseq), the high cost and the access to large computational resources often result in expensive, time-consuming analysis of a limited number of RNA samples. However, RT-QPCR gene expression is relatively easy to perform, even by non-expert users and constitute a fast and accessible method to access plant transcriptome (Fassbinder-Orth, 2014). Here we report a new database of key response genes of rice involved in different major processes in leaves. Thanks to the Fluidigm technology, we access for the first time to the simultaneous transcriptomics regulation of 88 marker genes under 9 different physiological situations.

Overall, our study confirms a majority of the regulation data found in the bibliography (~ 64%) (**Figure 2**; **Supplementary Table.S3)**. For example, the gene family *OsJAZ* (*OsJAZ5*, *OsJAZ6*, *OsJAZ11*) was strongly upregulated by the MeJA treatment and by drought, as expected (Dhakarey et al., 2017; Riemann et al., 2015). Similarly, a high number of genes were both upregulated by drought and ABA, which is not surprising because ABA is a part of the drought stress response in rice (Cai et al., 2015). However we also discovered non expected new regulations; for instance, ABA responsive genes like *ABI2* and *RCCR1* were induced in ABA treatment but also by the kinetin treatment, an unexpected regulation since ABA and CK are usually play antagonist roles in rice leaves (Zhang et al., 2021). Consequently, from all the genes tested, only 8 were specifically responding to the treatment they were originally described to be responsive. This result is not surprising since a large majority of available studies only analyzed the regulation of genes in one condition without checking possible other regulations (Biswas et al., 2017).

Principal component analysis of the expression data (PCA, **Figure 4**) revealed an anticorrelation of auxin and cytokinin treatments on rice leaf gene expression, a well-known result in rice and Arabidopsis (De Vleesschauwer et al., 2014, 2013). PCA also suggested that ABA treatment and nitrogen depletion produced anti-correlated transcriptomic responses, which is consistent with recent results showing that nitrogen stress induces reduction of ABA responsive genes in rice (Zakari et al., 2020). However it was not excepted to see that ABA treatment and *M. oryzae* infection produced opposite patterns of gene expression; indeed *M. oryzae* infection is associated with elevated ABA levels and reprogramming of ABA responsive gene by producing its own ABA to hijack rice defense (Ribot et al., 2008; Takatsuji and Jiang, 2014). Since SA, JA and ET treatments can mimic rice infections by different pathogens (Morimoto et al., 2018; Nahar et al., 2011), we expected to see some correlated transcriptomic responses with inoculation with *M. oryzae*. Surprisingly the defense hormones SA and JA appeared to produce anti-correlated expression patterns whereas ET pattern correlated with the pattern induced by inoculation of *M. oryzae*. This result can be partially explained by the young age of the plant since SA and JA activation of resistance depends on the age of the plant and mostly occurs in adult plant (De Vleesschauwer et al., 2013; Iwai et al., 2007) compared to ET which can mediate resistance even in rice seedling (Nahar et al., 2011). Altogether, these observations plead for a better and integrated comparison of pathogen defense and JA/SA/ET/ABA responses.

**Figure 4.**
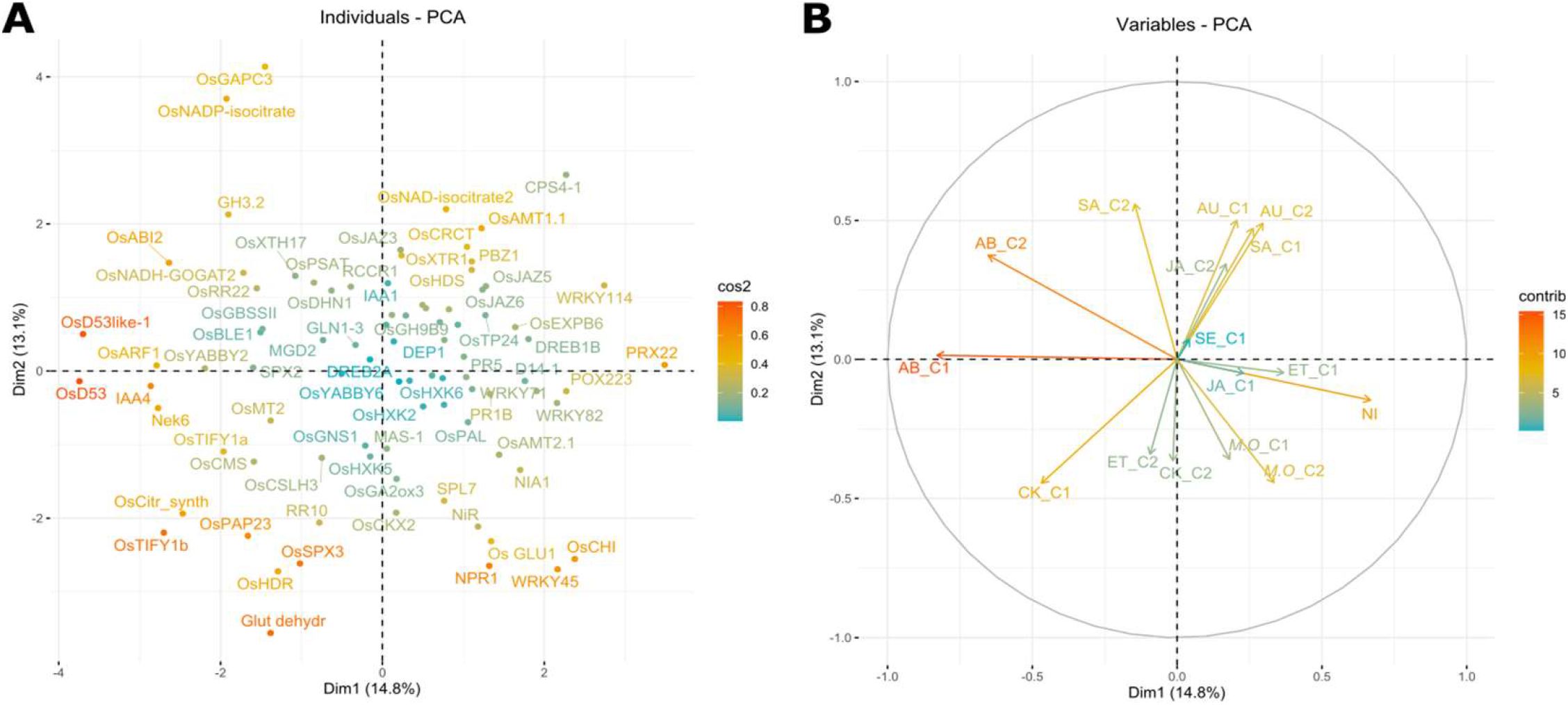
Principal component Analysis (PCA) of transcriptomic response of 88 marker genes in nine treatments. Gene targeted were set as variable and condition (treatment / concentration) as individual for PCA analysis. Data are visualized on a (A) Principal component analysis of the gene expression (fold change compared to mock treatments) of all genes. PCA has been done on the 88 marker genes and the color represents their “cos2” values. (B) Variable projection color by their contribution to axis. For most treatments, two doses were available and are shown (e.g. SA_C1 and SA_C2).

In conclusion, we gathered 88 genes transcriptionally regulated during key rice leaf processes. Monitoring their transcriptional regulation under different treatments allowed us to confirm previous regulation known in the bibliography and to discover new gene regulations. Additionally, this pool of genes represents a unique database for researchers to rapidly access and explore at low cost rice leaf transcriptome with Fluidigm technology or with classical RT-QPCR analysis. For future experience, the available gene expression dataset produced here can be used for any transcriptomic analysis on rice leaves to characterize the impact of unknown stresses by comparing it to our treatments.

## Supplementary material

**Supplementary figure S1.**
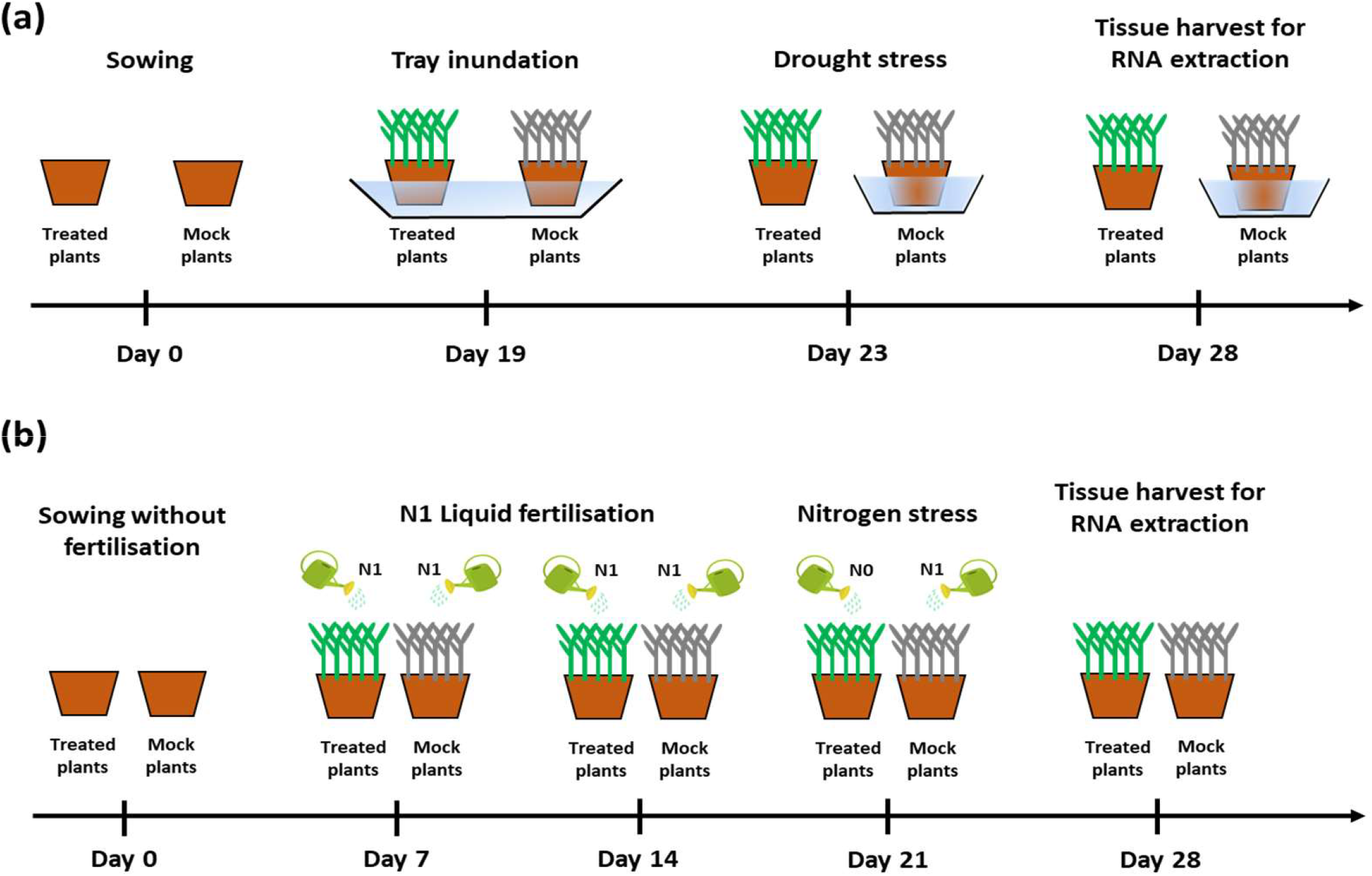
Experimental protocol used for drought and nitrogen treatment in this study. Each treatment was performed on at least 6 pots of 5 plants.

**Supplementary figure S2.**
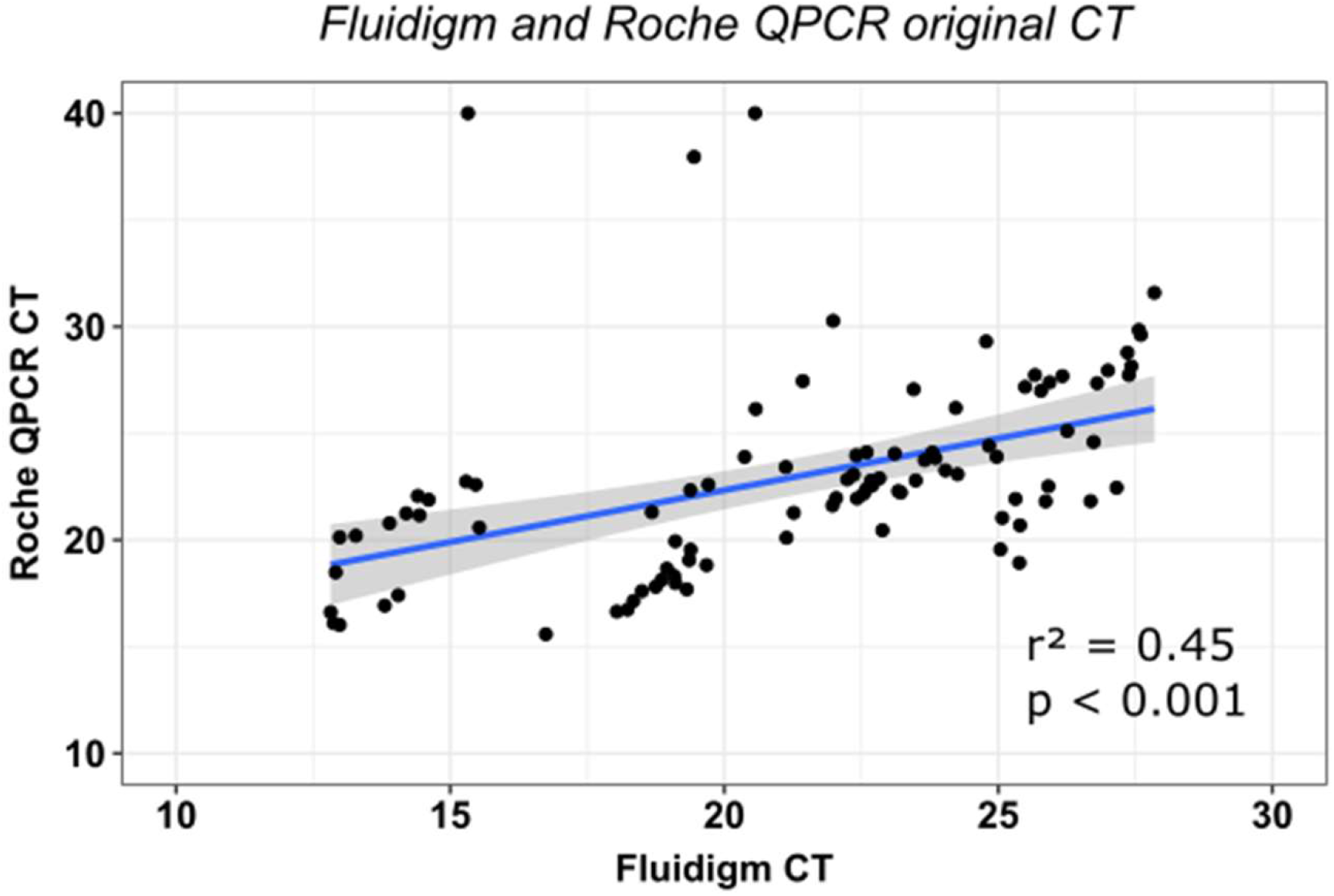
Relation between Fluidigm and QPCR values on the same samples used in this study. Data were selected randomly in the 9216 values obtain from Fluidigm analysis. The corresponding samples (pre-amplified sample used from Fluidigm analysis) were subjected to classical Q-PCR analysis (see methods). Person correlation were calculated and R^2^ and p.value are shown.

**Supplementary Table.S1 Details of phytohormones and mock treatment used in this study.**

**Supplementary Table.S2A List of the genes selected, transcriptional regulation, process targeted, primers and reference based on bibliographic analysis**

Complete references cited in the Excel file are available in the word document **Table.S2B**

**Supplementary Table.S3. Table of fold changes**

Fold change were calculated by dividing gene expression in treated plant by expression in the corresponding Mock condition. Bold values indicate statistically different fold change from the other detected by Tukey HSD test (see methods).

